# Measuring “Intolerance to Mutation” in Human Genetics

**DOI:** 10.1101/382481

**Authors:** Zachary L. Fuller, Jeremy J. Berg, Hakhamanesh Mostafavi, Guy Sella, Molly Przeworski

## Abstract

In numerous applications, from working with animal models to mapping the genetic basis of human disease susceptibility, it is useful to know whether a single disrupting mutation in a gene is likely to be deleterious^1–4^. With this goal in mind, a number of measures have been developed to identify genes in which protein-truncating variants (PTVs), or other types of mutations, are absent or kept at very low frequency in large population samples—genes that appear “intolerant to mutation”^3,5–9^. One measure in particular, pLI, has been widely adopted^7^. By contrasting the observed versus expected number of PTVs, it aims to classify genes into three categories, labelled “null”, “recessive” and “haploinsufficient”^7^. Such population genetic approaches can be useful in many applications. As we clarify, however, these measures reflect the strength of selection acting on heterozygotes, and not dominance for fitness or haploinsufficiency for other phenotypes.

---

Experimental biologists and human geneticists are often interested in whether a single disrupting mutation, be it a protein-truncating variant (PTV) or a missense mutation, is likely to have a phenotypic effect. A related question is whether such a mutation will lead to a reduction in fitness of its carrier. The relationship between these two questions, between effects on phenotypes and on fitness, is not straight-forward, with many potential paths from genotype to phenotype to fitness. For instance, a single mutation could lead to a severe clinical phenotype, indicating that the gene is haploinsufficient or that there is a gain of function, yet have small or negligible effects on fitness unless homozygous. As examples, in ELN and BRCA2, a single PTV leads to a severe but late onset disease while homozygote PTVs are lethal^10–13^; thus mutations in the genes are clearly haploinsufficient, but are they dominant with regard to fitness? Conversely, a mutation in a highly pleiotropic gene could have a very weak and potentially subclinical effect on any particular phenotype, yet cumulatively inflict a severe cost on fitness^14^.

Following common practice in human genetics (*e.g*.,^2^), we refer to genes in which a single loss of function mutation has a discernible phenotypic effect in heterozygotes as “haploinsufficient” (at least with regard to that phenotype). In turn, we describe genes in which a single disrupting mutation has an evolutionary fitness effect in heterozygotes as “dominant” (see Box 1). Although the term “dominance” is also used to refer to the effect of a single allele on phenotype, for clarity, here, we restrict its use to denote effects on fitness. More precisely, following the convention in population genetics, we denote the fitnesses 1, 1 – *hs*, and 1 – *s* as corresponding, respectively, to genotypes AA, AD, and DD, where D is the deleterious allele, *h* is the dominance coefficient, and *s* is the selection coefficient. Thus, a mutation is completely recessive if *h* is equal to 0, that is if deleterious fitness effects are only present in homozygotes, and at least partially dominant otherwise.

Estimating the strength of selection acting on a gene in terms of the selection coefficient (*s*) and dominance effects (*h*) of mutations, has a long tradition in population genetics^15–18^. In model organisms, such estimates have relied on mutation accumulation experiments and assays of gene deletion libraries^15,19–21^; in humans and other species, these parameters have been inferred from patterns of genetic variation^22–26^. The inferences are based on the notion of a “mutation-selection-drift balance”, namely that the frequencies of deleterious alleles in a sample reflect a balance between the rate at which they are introduced by mutation and the rate at which they are purged from the population by selection (as well as change in frequency randomly due to genetic drift). Mutations with larger hs are purged more effectively and hence are expected to be at lower frequencies in the population—or, equivalently, are more likely to be absent from large samples (Box 1). Therefore, one way to identify genes whose loss is likely to reduce fitness is to assess whether disrupting mutations are found at lower frequencies than expected under some sensible null model.

## Box 1

Deleterious alleles are introduced into the population by mutation, then change in frequency due to the combined effects of genetic drift and natural selection. Unless a disease mutation confers an advantage in some environments (*e.g*., the sickle cell allele in populations with severe malaria^27^), the frequency at which it will be found in a population reflects a balance between the rate at which it is introduced by mutation and removed purifying selection, modulated by the effects of genetic drift ^28–30^.

This phenomenon is referred to as “mutation-selection-drift” balance and modeled as follows (*e.g*., see^31^). Let *u* be the mutation rate from the wild type allele A to deleterious allele D. This mutation rate can be defined per site or per gene, by summing the mutation rate to deleterious alleles across sites (this simple summing implicitly assumes that there is no complementation and compound heterozygotes for deleterious alleles have the same fitness effects as homozygotes^32^). The fitness of diploid individuals carrying genes with wild-type (A) or deleterious (D) alleles is given by

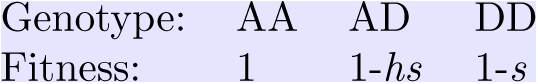

where *s* is the selection coefficient, which measures the fitness of DD relative to AA, and *h* is the dominance coefficient, such that *hs* is the reduction in fitness of AD relative to AA. In population genetics, the term dominance (with respect to fitness) is often defined as *h* > 0.5. Here, however, we define a mutation as partially dominant so long as *h* is not near 0, as this criterion is directly relevant to the expected frequency of deleterious mutations^33^.

In the limit of an infinite, panmictic population (*i.e*., ignoring genetic drift and inbreeding), when *h* > 0 (and *hs* ≫ *u*), the equilibrium frequency of the deleterious (D) allele, *q*, is approximately^29^:

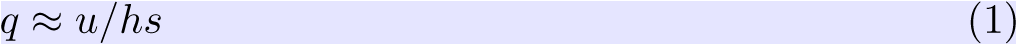

Notably, when *h* > 0, the equilibrium frequency *q* is determined by the strength of selection in heterozygotes (*i.e., hs*, the joint effects of *h* and *s*) because deleterious homozygotes are too infrequent for selection on them to have an appreciable effect on allele dynamics in the population. Hence, in this approximation, for a given *hs*, different combinations of *h* and *s* will yield the same frequency of *q*.

Under the same conditions, for a completely recessive allele (*h* = 0), *q* is well approximated by^29^:

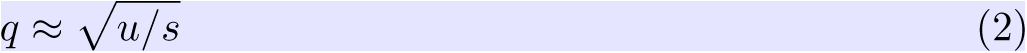

Here, the equilibrium frequency is determined by selection in homozygotes. In this limit of an infinite population size, the frequency corresponding to a recessive allele with a given *s* > 0 can also arise from a dominant allele for some value of *hs* > 0.

In a finite population, there is a distribution of deleterious allele frequencies rather than a single (deterministic) value for any values of *h* and *s*. For a constant population size *N*, this distribution was derived by Wright^30^ and is again a function of *hs* (assuming that *2Nhs* ≫ 1 and setting aside the case of sustained, high levels of inbreeding^34^). The resulting distribution can be highly variable, reflecting both stochasticity in the mutation process and the variance due to genetic drift. Dramatic changes in population size, as experienced by human populations, can also have a marked effect on the distribution of deleterious alleles. Regardless of these complications, it remains the case that distinguishing complete recessivity (*h* = 0) from small *hs* may not be feasible and that, other than for complete recessivity, the expected allele frequency is a function of *hs*, not *h* and *s* separately^33^.

To our knowledge, this approach—of prioritizing human disease genes on the basis of fitness consequences of disrupting mutations—was introduced by Petrovski et al.^5^, who ranked genes by comparing the observed number of common PTVs and missense mutations to the total number of observed variants. Their statistic was then supplemented by a number of others^6,35–37^, notably pLI, which is defined as an estimate of the “probability of being loss of function intolerant”^7^. Loosely, pLI is derived from a comparison of the observed number of PTVs in a sample to the number expected in the absence of fitness effects (*i.e*., under neutrality), given an estimated mutation rate for the gene. Specifically, Lek et al.^7^ assumed that the number of PTVs observed in a gene is Poisson distributed with mean λ*M*, where *M* is the number of segregating PTVs expected in a sample under neutrality (estimated for each gene based on a mutation model^6^ and the observed synonymous polymorphism counts) and λ reflects the depletion in the number due to selection. The authors considered that a gene can be “neutral” (with λ_*Null*_ = 1), “recessive” (λ_*Rec*_ = 0.463) or “haploinsufficient” (λ_*HI*_ = 0.089). The fixed values of λ_*Rec*_ and λ_*HI*_ were obtained from the average reduction in the number of observed PTVs in genes classified as “recessive” and “severe haploinsufficient”, respectively; the classification was based on phenotypic effects of mutations in the ClinGen dosage sensitivity gene list and a hand curated gene set of Mendelian disorders^38^. Given this model, Lek et al.^7^ estimated the proportion of human genes in each of their three categories and then, for any given gene, they obtained the maximum *a posteriori* probability that it belongs to each of the categories. Genes with high probability (set at ≥ 0.9) of belonging to the “haploinsufficient” class were classified as “extremely loss of function intolerant” ^7^.

pLI has been broadly used in human genetics, to help identify genes in which a single disrupting mutation is likely of clinical significance^4,39-46^. It is also increasingly employed in clinical annotation and in databases of mouse models, as indicative of haploinsufficiency and dosage sensitivity^47-51^. In fact, however, pLI and related measures reflect only the strength of selection acting on heterozygotes and are not directly informative about dominance effects on fitness, let alone about the degree of haploinsufficiency with respect to a phenotype.

The reason can be understood in population genetic terms: unless *h* is vanishingly small (or long-term inbreeding levels are very high), a reduction in the frequency of PTVs—and hence of PTV counts—is indicative of the strength of selection acting on heterozygotes, *hs*, and not of the two parameters *h* and *s* separately. This result derives from mutation-selection-drift balance theory developed by Haldane^28,29^, Wright^30^, and others^52^ (see Box 1). Intuitively, it reflects the fact that when fitness effects in heterozygotes are strong relative to the rate of genetic drift, deleterious alleles are kept at low frequency in the population. Homozygotes for the deleterious allele are therefore exceedingly rare and selection acts almost entirely through heterozygotes. As a result, the frequencies of PTVs in a sample—and therefore pLI and related measures—reflect the strength of selection acting on heterozygotes. This may be true even for genes classified as phenotypically recessive by clinicians: although a much stronger phenotype is seen in homozygotes, a subtle fitness effect on heterozygotes can be sufficient to markedly decrease the frequency of disease mutations^53^.

To illustrate this point, we used forward simulations to model how the observed counts of PTVs (and hence pLI) depend on *h* and *s* for a gene of typical length, considering both a constant size population setting (Fig 1A, see legend for details) and a more realistic model for human demographic history^54^(Fig 1B).

**Figure 1:**
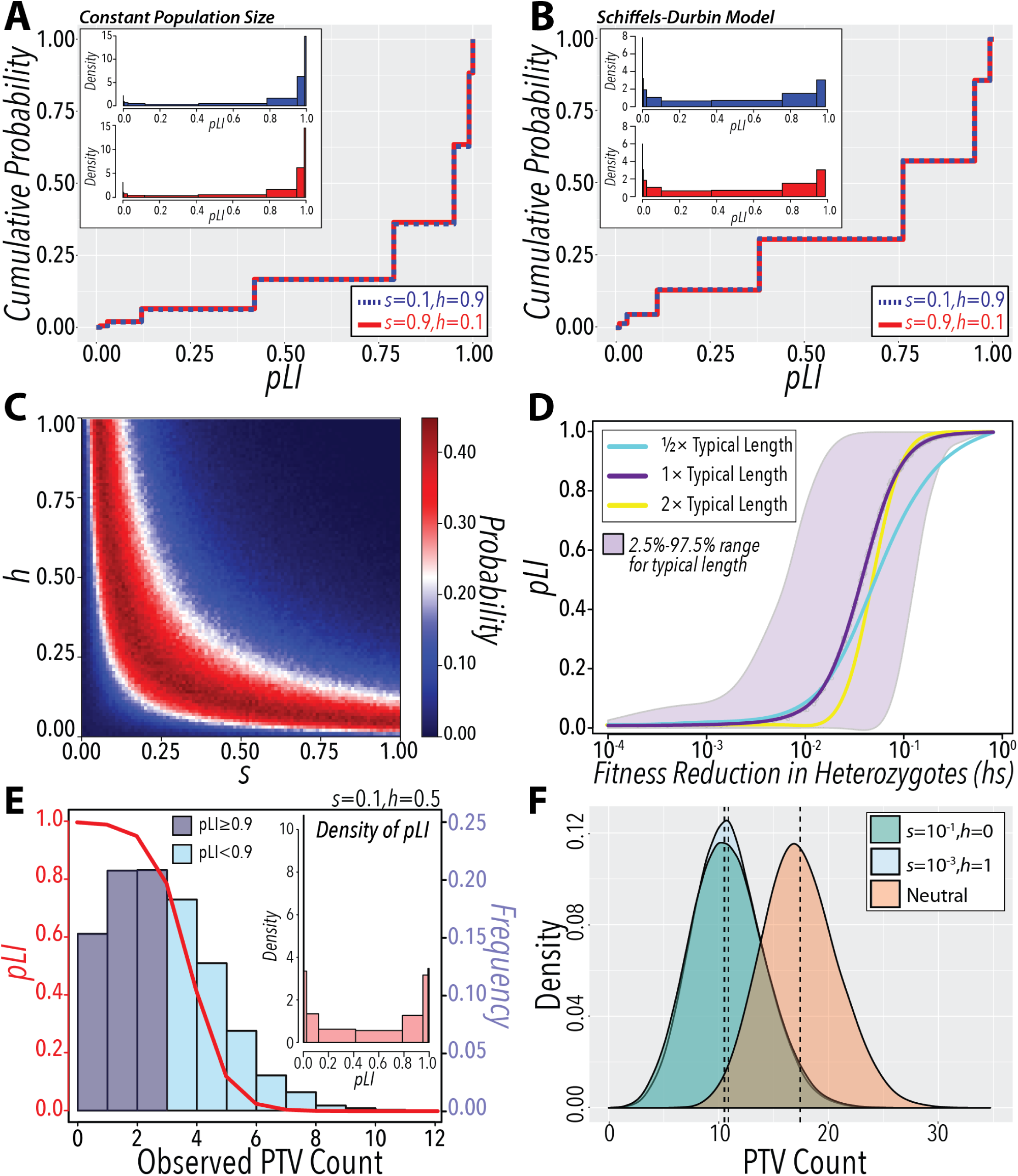
Properties of pLI. **(A & B) Different combinations of *h* and *s* with the same *hs* value yield highly similar distributions of pLI**. We considered PTVs arising in a hypothetical human gene of typical length (*i.e*., 225 PTV mutational opportunities, the average number in the human genome) for (A) a population of constant size and (B) a plausible model of changes in the effective population size of Europeans^54^. Mutations arise at rate *u* = 1.5 × 10^−8^ per mutational opportunity. While this value of *u* is only approximate, it yields realistic numbers of PTVs; the qualitative conclusions are the same for other choices. Subsequent generations are formed by Wright-Fisher sampling with selection, modeled by choosing parents for each generation according to their fitness. We assumed no intragenic recombination and that a PTV mutation can only occur on a background free of other PTV mutations (as is highly likely). In the constant population model, we set *N* = 100,000, reflective of the more recent time period relevant to the dynamics of deleterious mutations^53^, and ran each simulation for 10*N* generations. The number of segregating PTVs are estimated from a sample size of 33,370 diploid individuals drawn at present, to match the number of non-Finnish Europeans in ExAC^7^. In the more realistic demographic model for Europeans^54^, each simulation begins with a constant population size *N* of 14,448 (the ancestral size inferred by^54^) and a burn-in of 10*N* generations, before the first population size change 55,940 generations ago (following^14^). We obtained the mean number of PTVs under neutrality, *i.e*., for *h* = 0 and *s* = 0, by averaging over 10^6^ simulations. We then ran 10^6^ replicates for each combination of *s* and *h*, recording the distinct number of PTVs that are segregating at present. For each replicate, we calculated the ratio λ of the observed count of distinct segregating PTVs to the expected number under neutrality. Following the procedure in^7^, we then calculated pLI using the observed λ and the estimates of the mixing weights for each set obtained from ExAC (*π*_*Null*_ = 0.208, *π*_*Rec*_ = 0.489, and *π*_*HI*_ = 0.304). The insets in each figure show the density of the distribution of pLI scores. Since we used the true expected number of PTVs under neutrality, rather than an estimate (as is the case in practice^7^), we are somewhat under-estimating the variability in pLI scores. **C++** code and detailed methods for the simulations are available at https://github.com/zfuller5280/MutationIntoleranceSimulations. **(C) The probability of observing a specific PTV count is maximized along a ridge of fixed *hs***. The figure depicts the probability of observing the PTV count for a gene with the same mutational parameters as Fig 1A, and *s* = 0.10, *h* = 0.90, and assuming a plausible model of population size changes^54^ (see the legend of Fig 1A). The first simulation replicate happened to yield 3 PTVs, which we used as our “observed” PTV count. We estimated the likelihood of *h* and *s*, *i.e*., the probability of this “observed” value, for a grid of *h* and *s* values, using 10^6^ replicates for each parameter combination. **(D) Behavior of pLI as a function of *hs***. We simulated the counts of PTVs under a plausible model of population size changes in Europeans^54^, for a range of *hs* values (see the legend of Fig 1A). For each run, we calculated pLI using the observed number of PTVs and the expected number obtained from averaging over 10^6^ simulations with *h* = 0 and *s* = 0. The gray circles depict the average pLI over 10^6^ simulations for each value of *hs*, shown on the x-axis (on a log_10_ scale), in a human gene of typical length; the dark purple line is the loess smoothed curve over all simulations. The shaded area represents the 2.5^th^ and 97.5^th^ percentiles of pLI scores for each value of *hs*. The cyan and yellow lines are the loess smoothed curves for simulations in a human gene with half and twice the number of PTV mutational opportunities, respectively. **(E) For a given *hs*, pLI scores are highly variable**. Considering *s* = 0.1, *h* = 0.5, we generated the distribution of pLI scores in a human gene of typical length, with the expected number of PTVs under neutrality obtained as described in the legend of Fig 1A. The red curve depicts the pLI score as a function of the number of observed PTVs (calculated as in^7^). The histogram represents the distribution of simulated PTV counts under a plausible model of historical population size changes in Europeans^54^ (see the legend of Fig 1A), with darker shaded bars indicating pLI values that would be classified as “extremely loss-of-function intolerant” ^7^. The inset shows the density of pLI scores. **(F) Complete recessivity (*h*=0) can lead to similar PTV counts as weak selection on heterozygotes (*hs*>0)**. We simulated the counts of PTVs in a human gene of typical length undera plausible model of population size changes in Europeans^54^, for different combinations of *h* and *s* (see legends of Fig 1A). The distribution labeled as neutral depicts the counts of PTVs in simulations with *h* and *s* both equal to 0. Each distribution shows the results from 10^6^ simulations. Dashed lines indicate the mean for each distribution.

As can be seen, markedly different combinations of *h* and *s* lead to indistinguishable distributions of PTV counts (and hence of pLI values), so long as *hs* is the same (Fig 1A, B). More generally, the probability of observing a specific PTV count is maximized along a ridge corresponding to combinations of *h* and *s* that result in a given *hs* value (Fig 1C). As a result, pLI can be near 1 even when the dominance coefficient *h* is small, provided *s* is sufficiently large, and is therefore not indicative of dominance *per se*.

Although these considerations make clear that pLI should be thought of as reflecting *hs*, it was not designed to be an estimator of this parameter, and has several problematic features as such. First, for a given value of *hs*, the expected value of pLI varies with gene length (Fig 1D). Second, for a typical gene length and a wide range of *hs* values (*i.e*., 10^−3^-10^−1^), the distribution of pLI is highly variable and bimodal, covering most of the range from 0 to 1 (Fig 1E). Consequently, two genes with the same *hs* can be assigned radically different pLI values (Fig 1E). Conversely, the same pLI value can reflect markedly different *hs* values, as illustrated by the large variance of pLI in the *hs* range between 10^−3^ and 10^−1^ (Fig 1D). Outside this range of *hs* values, pLI is almost uninformative about the underlying parameter: below *hs* ≈ 10^−3^, pLI is ~ 0 for any value of *hs;* above *hs* ≈ 10^−1^, it is always ~ 1, properties that worsen with increasing gene length (Fig 1D). Our simulations further illustrate that for a given *hs*, genetic drift also contributes to the variance in PTV counts, a feature that is ignored in the construction of pLI (through its reliance on a Poisson distribution of PTV counts)^55^. Thus, if the goal is to learn about fitness effects to help prioritize disease genes, a direct estimate of *hs* (*e.g*.,^9,55^) under a plausible demographic model, together with a measure of statistical uncertainty, would be preferable.

Recasting pLI in a population genetic framework also helps to understand why the “recessive” assignments are even less reliable^7^. Lek et al.^7^ aim to divide genes into three categories, two of which correspond to *hs* > 0 (pLI) and *hs* = 0, *s* = 0 (pNULL). Logically, the remaining category (pREC) should include completely recessive cases (*i.e*., where *hs* = 0 but *s* > 0), in which selection acts exclusively against homozygotes (Box 1). Regardless of the method used, however, it can be infeasible to distinguish this category from the *hs* > 0 case, because the same expected allele frequency (and hence PTV count) can arise when *h* = 0 or when *hs* > 0 but small (see Box 1 and Fig 1F). For example, for a typical per gene mutation rate to disease alleles of *u* = 10^−6^ and no genetic drift, the frequency of disease alleles would be 1% whether *h* = 0 (completely recessive) and *s* = 10^−2^ or *h* = 1 (fully dominant) and *s* = 10 (see equations in Box 1). In other words, strongly deleterious, completely recessive PTVs can be hard to distinguish from those that are weakly selected and at least partially dominant.

Why then, in practice, are genes classified by clinicians as “dominant” based on Mendelian disease phe-notypes enriched for high pLI scores compared to those classified as “recessive”^4,7,41^? Mendelian disease genes consist mostly of cases in which mutations are known to cause a highly deleterious outcome, *i.e*., for which there is prior knowledge that *s* is likely to be large (even close to 1). When *s* is large, a gene will be classified by pLI as haploinsufficient so long as fitness effects in heterozygotes are sufficient to decrease the number of observed PTVs, *i.e*., so long as *h* is not tiny. For most genes, however, there is no prior knowledge about *s*, and in that case, pLI—or any measure based on the frequency of PTVs—cannot reliably distinguish recessivity from dominance, let alone identify haploinsuffi-ciency.

In summary, population genetic approaches based on the deficiency of putatively deleterious mutations^3,6,9,36,56^ hold great promise for prioritizing genes in which mutations are likely to be harmful in heterozygotes^5,9^. Recasting these approaches in terms of underlying population genetic parameters provides a natural framework for their interpretation and a clearer understanding of what they can reliably infer: these approaches identify genes in which single PTVs likely have large fitness effects in heterozygotes. For this subset of genes, there is information about dominance when s is known to be large and not otherwise. Moreover, for no genes can the methods be used to infer haploinsufficiency status. Where fitness effects are to be used as an indication of pathogenicity, we therefore argue that a better approach is the development of direct estimates of *hs* (and measures of uncertainty) under realistic demographic models for the population of interest.

## Acknowledgments

We thank A. Chakravarti, G. Coop, M.B. Eisen, M. Hurles, J.K. Pritchard, and Y. Shen for helpful discussions. This work was supported by GM128318 to ZF, GM126787 to JB, GM121372 to MP and GM115889 to GS. We acknowledge computing resources from Columbia University’s Shared Research Computing Facility project, which is supported by NIH Research Facility Improvement Grant 1G20RR030893-01, and associated funds from the New York State Empire State Development, Division of Science Technology and Innovation (NYSTAR) Contract C090171. Code used to run forward simulations of PTV counts is available on-line

## References

[1] J. A. Blake, C. J. Bult, J. T. Eppig, J. A. Kadin, and J. E. Richardson, The Mouse Genome Database genotypes∷phenotypes, Nucleic Acids Research 37, D712 (2009), ISSN 0305-1048.

[2] N. Huang, I. Lee, E. M. Marcotte, and M. E. Hurles, Characterising and Predicting Haploinsufficiency in the Human Genome, PLOS Genetics 6, e1001154 (2010), ISSN 1553-7404.

[3] K. Eilbeck, A. Quinlan, and M. Yandell, Settling the score: variant prioritization and Mendelian disease, Nature Reviews. Genetics 18, 599 (2017), ISSN 1471-0064.

[4] I. Bartha, J. d. Iulio, J. C. Venter, and A. Telenti, Human gene essentiality, Nature Reviews Genetics 19, 51 (2018), ISSN 1471-0064.

[5] S. Petrovski, Q. Wang, E. L. Heinzen, A. S. Allen, and D. B. Goldstein, Genic Intolerance to Functional Variation and the Interpretation of Personal Genomes, PLOS Genetics 9, e1003709 (2013), ISSN 1553-7404.

[6] K. E. Samocha, E. B. Robinson, S. J. Sanders, C. Stevens, A. Sabo, L. M. McGrath, J. A. Kosmicki, K. Rehnstrm, S. Mallick, A. Kirby, et al., A framework for the interpretation of de novo mutation in human disease, Nature Genetics 46, 944 (2014), ISSN 1061-4036.

[7] M. Lek, K. J. Karczewski, E. V. Minikel, K. E. Samocha, E. Banks, T. Fennell, A. H. ODonnell-Luria, J. S. Ware, A. J. Hill, B. B. Cummings, et al., Analysis of protein-coding genetic variation in 60,706 humans, Nature 536, 285 (2016), ISSN 0028-0836.

[8] Y.-F. Huang, B. Gulko, and A. Siepel, Fast, scalable prediction of deleterious noncoding variants from functional and population genomic data, Nature Genetics 49, 618 (2017), ISSN 1546-1718.

[9] C. A. Cassa, D. Weghorn, D. J. Balick, D. M. Jordan, D. Nusinow, K. E. Samocha, A. O’Donnell-Luria, D. G. MacArthur, M. J. Daly, D. R. Beier, et al., Estimating the selective effects of heterozygous protein-truncating variants from human exome data, Nature Genetics 49, 806 (2017), ISSN 1061-4036.

[10] M. C. Raybould, A. J. Birley, and M. Hultn, Molecular variation of the human elastin (ELN) gene in a normal human population, Annals of Human Genetics 59, 149 (1995), ISSN 1469-1809.

[11] R. Wooster, G. Bignell, J. Lancaster, S. Swift, S. Seal, J. Mangion, N. Collins, S. Gregory, C. Gumbs, G. Micklem, et al., Identification of the breast cancer susceptibility gene BRCA2, Nature 378, 789 (1995), ISSN 1476-4687.

[12] J. E. Wagenseil, C. H. Ciliberto, R. H. Knut-sen, M. A. Levy, A. Kovacs, and R. P. Mecham, The importance of elastin to aortic development in mice, American Journal of Physiology - Heart and Circulatory Physiology 299, H257 (2010), ISSN 0363-6135.

[13] R. Roy, J. Chun, and S. N. Powell, BRCA1 and BRCA2: different roles in a common pathway of genome protection, Nature Reviews. Cancer 12, 68 (2011), ISSN 1474-1768.

[14] Y. B. Simons, K. Bullaughey, R. R. Hudson, and G. Sella, A population genetic interpretation of GWAS findings for human quantitative traits, PLOS Biology 16, e2002985 (2018), ISSN 1545-7885.

[15] M. J. Simmons and J. F. Crow, Mutations affecting fitness in Drosophila populations, Annual Review of Genetics 11, 49 (1977), ISSN 0066-4197.

[16] P. D. Keightley, The distribution of mutation effects on viability in Drosophila melanogaster, Genetics 138, 1315 (1994), ISSN 0016-6731.

[17] H. W. Deng and M. Lynch, Estimation of Deleterious-Mutation Parameters in Natural Populations, Genetics 144, 349 (1996), ISSN 0016-6731.

[18] H. A. Orr, Fitness and its role in evolutionary genetics, Nature reviews. Genetics 10, 531 (2009), ISSN 1471-0056.

[19] T. Mukai, S. I. Chigusa, L. E. Mettler, and J. F. Crow, Mutation Rate and Dominance of Genes Affecting Viability in DROSOPHILA MELANOGASTER, Genetics 72, 335 (1972), ISSN 0016-6731.

[20] N. Phadnis and J. D. Fry, Widespread Correlations Between Dominance and Homozygous Effects of Mutations: Implications for Theories of Dominance, Genetics 171, 385 (2005), ISSN 0016-6731.

[21] A. F. Agrawal and M. C. Whitlock, Inferences About the Distribution of Dominance Drawn From Yeast Gene Knockout Data, Genetics 187, 553 (2011), ISSN 0016-6731, 1943-2631.

[22] S. H. Williamson, R. Hernandez, A. Fledel-Alon, L. Zhu, R. Nielsen, and C. D. Bustamante, Simultaneous inference of selection and population growth from patterns of variation in the human genome, Proceedings of the National Academy of Sciences of the United States of America 102, 7882 (2005), ISSN 0027-8424.

[23] A. Eyre-Walker, M. Woolfit, and T. Phelps, The Distribution of Fitness Effects of New Deleterious Amino Acid Mutations in Humans, Genetics 173, 891 (2006), ISSN 0016-6731.

[24] A. R. Boyko, S. H. Williamson, A. R. Indap, J. D. Degenhardt, R. D. Hernandez, K. E. Lohmueller, M. D. Adams, S. Schmidt, J. J. Sninsky, S. R. Sunyaev, et al., Assessing the Evolutionary Impact of Amino Acid Mutations in the Human Genome, PLOS Genetics 4, e1000083 (2008), ISSN 1553-7404.

[25] F. Racimo and J. G. Schraiber, Approximation to the Distribution of Fitness Effects across Functional Categories in Human Segregating Polymorphisms, PLOS Genetics 10, e1004697 (2014), ISSN 1553-7404.

[26] B. Y. Kim, C. D. Huber, and K. E. Lohmueller, Inference of the Distribution of Selection Coefficients for New Nonsynonymous Mutations Using Large Samples, Genetics 206, 345 (2017), ISSN 1943-2631.

[27] F. B. Piel, A. P. Patil, R. E. Howes, O. A. Nyangiri, P. W. Gething, T. N. Williams, D. J. Weatherall, and S. I. Hay, Global distribution of the sickle cell gene and geographical confirmation of the malaria hypothesis, Nature Communications 1, 104 (2010), ISSN 2041-1723.

[28] J. B. S. Haldane, A Mathematical Theory of Natural and Artificial Selection, Part V: Selection and Mutation, Mathematical Proceedings of the Cambridge Philosophical Society 23, 838 (1927), ISSN 1469-8064, 0305-0041.

[29] J. B. S. Haldane, The Effect of Variation of Fitness, The American Naturalist 71, 337 (1937), ISSN 0003-0147.

[30] S. Wright, The Distribution of Gene Frequencies in Populations, Proceedings of the National Academy of Sciences of the United States of America 23, 307 (1937), ISSN 0027-8424.

[31] J. H. Gillespie, Population Genetics: A Concise Guide (JHU Press, 2004), ISBN 978-0-8018-8008-7.

[32] A. G. Clark, Mutation-selection balance with multiple alleles, Genetica 102-103, 41 (1998), ISSN 0016-6707.

[33] Y. B. Simons, M. C. Turchin, J. K. Pritchard, and G. Sella, The deleterious mutation load is insensitive to recent population history, Nature Genetics 46, 220 (2014), ISSN 1061-4036.

[34] B. Charlesworth, Elements of Evolutionary Genetics (Roberts and Company Publishers, 2010), ISBN 978-0-9815194-2-5, google-Books-ID: dgN-FAQAAIAAJ.

[35] J. Steinberg, F. Honti, S. Meader, and C. Webber, Haploinsufficiency predictions without study bias, Nucleic Acids Research 43, e101 (2015), ISSN 0305-1048.

[36] I. Bartha, A. Rausell, P. J. McLaren, P. Mohammadi, M. Tardaguila, N. Chaturvedi, J. Fellay, and A. Telenti, The Characteristics of Heterozygous Protein Truncating Variants in the Human Genome, PLOS Computational Biology 11, e1004647 (2015), ISSN 1553-7358.

[37] J. Fadista, N. Oskolkov, O. Hansson, L. Groop, and J. Hancock, LoFtool: a gene intolerance score based on loss-of-function variants in 60 706 individuals, Bioinformatics 33, 471 (2017), ISSN 1367-4803.

[38] R. Blekhman, O. Man, L. Herrmann, A. R. Boyko, A. Indap, C. Kosiol, C. D. Bustamante, K. M. Teshima, and M. Przeworski, Natural selection on genes that underlie human disease susceptibility, Current biology: CB 18, 883 (2008), ISSN 0960-9822.

[39] S. H. Lelieveld, M. R. F. Reijnders, R. Pfundt, H. G. Yntema, E.-J. Kamsteeg, P. d. Vries, B. B. A. d. Vries, M. H. Willemsen, T. Kleefstra, K. Lhner, et al., Meta-analysis of 2,104 trios provides support for 10 new genes for intellectual disability, Nature Neuroscience 19, 1194 (2016), ISSN 1546-1726.

[40] D. M. Ruderfer, T. Hamamsy, M. Lek, K. J. Karczewski, D. Kavanagh, K. E. Samocha, Exome Aggregation Consortium, M. J. Daly, D. G. MacArthur, M. Fromer, et al., Patterns of genic intolerance of rare copy number variation in 59,898 human exomes, Nature Genetics 48, 1107 (2016), ISSN 1546-1718.

[41] J. A. Kosmicki, K. E. Samocha, D. P. Howrigan, S. J. Sanders, K. Slowikowski, M. Lek, K. J. Karczewski, D. J. Cutler, B. Devlin, K. Roeder, et al., Refining the role of de novo protein-truncating variants in neurodevelopmental disorders by using population reference samples, Nature Genetics 49, 504 (2017), ISSN 1546-1718.

[42] C. M. Skraban, C. F. Wells, P. Markose, M. T. Cho, A. I. Nesbitt, P. Y. B. Au, A. Begtrup, J. A. Bernat, L. M. Bird, K. Cao, et al., WDR26 Causes a Recognizable Syndrome of Intellectual Disability, Seizures, Abnormal Gait, and Distinctive Facial Features, The American Journal of Human Genetics 101, 139 (2017), ISSN 0002-9297.

[43] P. Stankiewicz, T. N. Khan, P. Szafranski, L. Slattery, H. Streff, F. Vetrini, J. A. Bernstein, C. W. Brown, J. A. Rosenfeld, S. Rednam, et al., Haploinsufficiency of the Chromatin Remodeler BPTF Causes Syndromic Developmental and Speech Delay, Postnatal Microcephaly, and Dysmorphic Features, The American Journal of Human Genetics 101, 503 (2017), ISSN 0002-9297.

[44] H. T. Nguyen, J. Bryois, A. Kim, A. Dobbyn, L. M. Huckins, A. B. Munoz-Manchado, D. M. Ruderfer, G. Genovese, M. Fromer, X. Xu, et al., Integrated Bayesian analysis of rare exonic variants to identify risk genes for schizophrenia and neurodevelopmental disorders, Genome Medicine 9, 114 (2017), ISSN 1756-994X.

[45] M. Zarrei, D. L. Fehlings, K. Mawjee, L. Switzer, B. Thiruvahindrapuram, S. Walker, D. Merico, G. Casallo, M. Uddin, J. R. MacDonald, et al., De novo and rare inherited copy-number variations in the hemiplegic form of cerebral palsy, Genetics in Medicine 20, 172 (2018), ISSN 1530-0366.

[46] H. O. Heyne, T. Singh, H. Stamberger, R. A. Jamra, H. Caglayan, D. Craiu, P. D. Jonghe, R. Guerrini, K. L. Helbig, B. P. C. Koeleman, et al., De novo variants in neurodevelopmental disorders with epilepsy, Nature Genetics 1 (2018), ISSN 1546-1718.

[47] M. Zech, S. Boesch, E. Maier, I. Borggraefe, K. Vill, F. Laccone, V. Pilshofer, A. Ceballos-Baumann, B. Alhaddad, R. Berutti, et al., Haploinsufficiency of KMT2b, Encoding the Lysine-Specific Histone Methyltransferase 2b, Results in Early-Onset Generalized Dystonia, American Journal of Human Genetics 99, 1377 (2016), ISSN 0002-9297.

[48] M. Haller, Q. Mo, A. Imamoto, and D. J. Lamb, Murine model indicates 22q11.2 signaling adaptor CRKL is a dosage-sensitive regulator of genitourinary development, Proceedings of the National Academy of Sciences 114, 4981 (2017), ISSN 0027-8424, 1091-6490.

[49] J. Wang, R. Al-Ouran, Y. Hu, S.-Y. Kim, Y.-W. Wan, M. F. Wangler, S. Yamamoto, H.-T. Chao, A. Comjean, S. E. Mohr, et al., MARRVEL: Integration of Human and Model Organism Genetic Resources to Facilitate Functional Annotation of the Human Genome, The American Journal of Human Genetics 100, 843 (2017), ISSN 0002-9297, 1537-6605.

[50] B. Afzali, J. Grnholm, J. Vandrovcova, C. OBrien, H.-W. Sun, I. Vanderleyden, F. P. Davis, A. Khoder, Y. Zhang, A. N. Hegazy, et al., BACH2 immunodeficiency illustrates an association between super-enhancers and haploinsufficiency, Nature immunology 18, 813 (2017), ISSN 1529-2908.

[51] N. Gosalia, A. N. Economides, F. E. Dewey, and S. Balasubramanian, MAPPIN: a method for annotating, predicting pathogenicity and mode of inheritance for nonsynonymous variants, Nucleic Acids Research 45, 10393 (2017), ISSN 0305-1048.

[52] J. F. Crow and M. Kimura, An introduction to population genetics theory (Harper & Row, 1970), google-Books-ID: ytMPAQAAMAAJ.

[53] C. E. G. Amorim, Z. Gao, Z. Baker, J. F. Diesel, Y. B. Simons, I. S. Haque, J. Pickrell, and M. Przeworski, The population genetics of human disease: The case of recessive, lethal mutations, PLOS Genetics 13, e1006915 (2017), ISSN 1553-7404.

[54] S. Schiffels and R. Durbin, Inferring human population size and separation history from multiple genome sequences, Nature genetics 46, 919 (2014), ISSN 1061-4036.

[55] D. Weghorn, D. J. Balick, C. Cassa, J. Kosmicki, M. J. Daly, D. R. Beier, and S. R. Sunyaev, Applicability of the mutation-selection balance model to population genetics of heterozygous protein-truncating variants in humans, bioRxiv 433961 (2018).

[56] J. M. Havrilla, B. S. Pedersen, R. M. Layer, and A. R. Quinlan, A map of constrained coding regions in the human genome., bioRxiv 220814 (2017).

